# 2-deoxy-D-glucose chemical exchange-sensitive spin-lock MRI of cerebral glucose metabolism after stroke in the rat

**DOI:** 10.1101/2025.01.08.628135

**Authors:** Philipp Boehm-Sturm, Patrick Schuenke, Marco Foddis, Susanne Mueller, Stefan P. Koch, Daniel J. Beard, Paul Holloway, Amin Mottahedin, Leif Schröder, Alastair M. Buchan, Philipp Mergenthaler

## Abstract

Rapid breakdown of cerebral glucose metabolism is a hallmark in stroke pathology. Metabolic activity delineates the penumbra from the infarct core, representing tissue that is potentially salvageable by therapeutic interventions. Tools to image dynamics of glucose and its spatial distribution could provide biomarkers of disease severity and of the success of therapeutic interventions.

Here, we developed a new protocol to measure glucose metabolism in a rat model of stroke using chemical exchange-sensitive spin-lock (CESL) MRI of the glucose analog 2-deoxy-D-glucose (2DG). We further implemented a protocol that combines 2DG-CESL-MRI with perfusion and diffusion MRI to relate this new signal to established definitions of hypoperfused tissue, cytotoxic edema and the penumbra. We found that 2DG-CESL-MRI provides a biomarker of disturbed glucose metabolism after stroke with high effect size. This is the first study to investigate CESL MRI of 2DG in the context of metabolism imaging in rodent stroke.

## Introduction

The brain almost exclusively relies on glucose as its main source of energy to sustain its function.^1, 2^ In ischemic stroke, occlusion of a cerebral blood vessel leads to a rapid decline of oxygen, glucose and other metabolites in the affected brain region. Acute neurodegeneration is triggered by a range of mechanisms including excitotoxicity, apoptosis, mitochondrial dysfunction, and neuroinflammation.^3-7^ Metabolic deficiency, and in particular impaired glucose metabolism, is an important mechanism controlling acute neurodegeneration in stroke.^3, 8^

The ischemic penumbra is defined as the hypoperfused brain tissue around the necrotic infarct core.^3, 9, 10^ The penumbra is characterized by decreased oxygen and glucose availability putting the cells of the neuro-glial-vascular unit in this tissue area at risk of death.^3^ However, if blood flow is restored, the penumbra is salvageable.^9, 10^ The penumbra is highly dynamic, with diverse cellular and molecular mechanisms influencing tissue outcomes.^3^ Clinical imaging of the penumbra typically focuses on blood flow and subsequent tissue damage.^11, 12^ Despite the early adoption of metabolic imaging in understanding the penumbra,^13-15^ it remains largely inaccessible in both preclinical and clinical settings. Therefore, developing scalable methods to measure glucose metabolism is crucial for improving diagnostic tools and therapeutic stratification in stroke. Established methods include ex vivo autoradiography of ^14^C-labeled glucose analogs such as 2-deoxy-D-glucose (2DG) and in vivo positron emission tomography (PET) of 18F 2-fluoro-2-desoxy glucose (18F-FDG).^16^ Both methods have their caveats: autoradiography is invasive and unsuitable for longitudinal studies, while 18F-FDG PET requires elaborate hardware and is heavily constrained by logistics such as proximity to a radiochemistry lab due to short half-life of the tracer and requires a dedicated radiation protection environment e.g. of nuclear medicine departments.

Magnetic resonance imaging (MRI) is broadly available and works with non-ionizing radiation. It has evolved as a key modality in stroke imaging since the contrast of MRI can be sensitized to different biological processes within one imaging session. For example, in ischemic stroke, dedicated sequences can measure reduced cerebral blood flow (CBF) and reduced apparent diffusion coefficient (ADC)^17^ to assess acute hypoperfusion and the extent of the lesion core, respectively. The mismatch of these signals provides a noninvasive marker of the penumbra.^15^ Hyperintense signal in T2-weighted (T2w) imaging provides a marker of vasogenic edema in subacute stroke and of necrotic tissue in older lesions.^18^ These findings have triggered MRI-based diagnostic concepts in the context of preclinical and clinical research and successfully translated to acute stroke patient care in many specialized stroke units.^18, 19^

Recently, MRI of natural D-glucose and glucose analogs has been achieved by utilizing chemical exchange between hydrogen bound to the molecule of interest and the larger water pool.^20-25^ Chemical exchange sensitive spin lock (CESL) MRI is a particularly signal-to-noise ratio (SNR) efficient technique that measures the relaxation rate 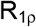 in the rotating frame.^26^ The goal of this study was to transfer these results to the context of stroke. We aimed to establish 2DG CESL MRI as a new biomarker of glucose metabolism in a rat model of ischemic stroke and combined it with T2w, CBF and ADC imaging to relate this new biomarker with established MRI markers of tissue damage.

## Material and Methods

### Phantom experiments

For the in vitro experiments, solutions of varying concentrations of 2DG in 1x Dulbecco’s phosphate buffered saline (DPBS, Gibco, Thermo Fisher Scientific, Hennigsdorf, Germany) were prepared and filled in 5 mm NMR tubes (Carl Roth GmbH, Karlsruhe, Germany). The pH in the phantoms was measured at ∼7.1. We note that the measurements are highly dependent on the exact buffer composition and pH.

### Animals and study approval

Male Wistar rats (∼8 weeks old, 280-320 g, n = 12) obtained from Charles River Germany were used in this study. Rats were housed in a temperature, humidity and light (12/12-h light/dark cycle; lights on: 06:00 am, experiments performed during the light cycle) controlled environment. All animal procedures were performed after approval by the regulating authority (Landesamt für Gesundheit und Soziales Berlin). Studies were performed in accordance with the German Animal Welfare Act and EU regulations, and reporting follows the ARRIVE guidelines.

### Anaesthesia and femoralis catheter

Animals were anesthetized using isoflurane in a 70%/30% N2O/O2 mixture. Anesthesia was induced with 5% isoflurane and maintained at 1.75-2.5% isoflurane during MCAO and 1.75% during MRI to achieve a breathing rate of ∼60-100/min. A catheter (Portex Fine Bore Polythene Tubing, 0.58 mm inner diameter, 0.96 mm outer diameter, 90 cm long) was surgically placed and secured in the femoral vein for 2DG i.v. injections during MRI measurements.

### Middle cerebral artery occlusion (MCAO) model of focal cerebral ischemia

Directly after placing the catheter, MCAO was surgically induced. A midline neck incision was made to expose the left common carotid artery, and a ligature was tied around the common carotid artery (CCA) and the external carotid artery (ECA). After a small incision into the CCA, a filament (390 µm diameter, 4039910PK10Re, Doccol, Sharon, MA/USA) was introduced into the CCA and advanced to the origin of the left MCA via the internal carotid artery. After 90 min, the filament was withdrawn. Both the CCA and ECA remained ligated during reperfusion. All MCAOs were performed by experienced surgeons. Blinding was not performed.

### MRI measurements

The experimental design is illustrated in Fig. 1. Directly after MCAO and reperfusion surgery, animals were transferred to a 7 T animal MRI (BioSpec 70/20 USR, Bruker, Ettlingen, Germany). Imaging was started within 10 to 15 min after reperfusion using an 86 mm diameter volume coil for transmission, a rat surface coil for reception and Paravision 6.0.1 software. An MR compatible monitoring system (SA Instruments, Stony Brook, NY, USA) was used to continuously control animal temperature via a rectal probe and respiration rate via a pressure sensitive pad under the thorax. A blanket connected to a heated circulating water system was used to keep the body temperature of the animal at 37.0 ± 0.5 °C. Animals were positioned prone on a dedicated animal holder inside the MRI, with ear and tooth bar fixation to minimize movement. First, a localizer scan was acquired followed by B0 mapping and second order shimming (MAPSHIM). A simultaneous mapping of water shift and B1 (WASABI) sequence was used to check B0 and B1 homogeneity.^27^ The protocol consisted of whole brain T2w MRI (2D RARE, 38 contiguous coronal slices with thickness 0.75 mm, field of view FOV = 33×33 mm^2^, image matrix MTX = 220×220, repetition time TR = 4000 ms, RARE factor 8, echo spacing 11 ms, effective echo time TE = 33 ms, bandwidth BW = 34.7 kHz, number of averages NA = 3, total acquisition time TA = 5:25 min), 2DG CESL MRI (single 2.25 mm thick slice with FOV and geometry matching three slices of the T2w through the rostral-caudal center of the striatum), MTX = 110×110, TR = 5.4 ms, TE = 2.4 ms, BW = 46.9 kHz, 13 spin lock times (TSL) between 5-200 ms, adiabatic tipping using optimized HSExp pulses^28^ and a matching spin lock pulse amplitude of B1 = 5 µT, NA = 1, 35 repetitions with TA = 1:26 min each, slow injection of 1 g/kg bodyweight 2DG i.v. after scan #5), diffusion MRI (SE-EPI, geometry matching the T2w, MTX = 128x128, TR = 3000 ms, TE = 20 ms, diffusion duration/separation = 2.5 ms/8.4 ms, 3 diffusion directions each with b = 650 s/mm^2^ and b = 1300 s/mm^2^, one b = 0 image, TA = 5:36 min), and arterial spin labeling MRI of CBF (FAIR-EPI, FOV = 33×33 mm^2^, MTX = 128x96, single 1.0 mm thick slice with geometry matching the CESL MRI, 16 inversion times between 35 ms-1600 ms, recovery time 10 s, TE = 16 ms, BW = 357.1 kHz, TA = 5:45 min). Perfusion MRI was performed using global shimming (not local MAPSHIM) to ensure sufficiently homogeneous B0 profile outside the FOV to enable efficient global inversion pulses. For phantom experiments, the same CESL scan was used but with slice thickness of 2 mm, FOV = 16x16 mm^2^, MTX = 128x128.

**Fig. 1:**
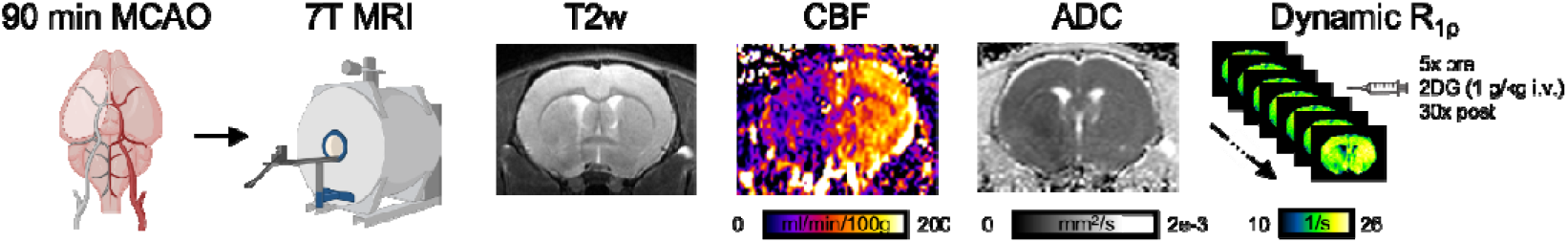
Study design. Rats underwent 90 min transient MCAO. After surgery, animals were directly transferred to a 7 T MRI system for T2-weighted (T2w) MRI, perfusion MRI of cerebral blood flow (CBF), diffusion MRI of apparent diffusion coefficient (ADC) followed by dynamic R1ρ mapping with CESL MRI before and after injection of 2DG.

### MRI data analysis

First, CESL images were motion-corrected in ANTx2 (https://github.com/ChariteExpMri/antx2)^29^ using rigid (6 degrees of freedom) image registration of the later (i = 2…35) to the first CESL image dataset. For each time point, the first spin lock time image (TSL = 5 ms) was used for registration, the transformation was then applied to the other 12 TSL images.

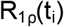 maps at time t_i_ were calculated voxel-wise via mono-exponential fitting of CESL image signal intensities S(t_i_) over spin lock time TSL using the CESL signal equation:

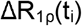 is a measure of local 2DG concentration and was calculated as the percent change with respect to the mean 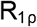 during the first five baseline CESL scans:

Region-of-interest (ROI) analysis was performed in two ways (Fig. 2) In the first approach, brain regions were automatically segmented on the different MRI contrasts (T2, ADC, CBF, CESL) using a series of 3D to 3D and 3D to 2D image registrations using the SIGMA rat brain template. The registration was then applied to the SIGMA atlas,^30^ which matches the template. For registration, we used the b = 0 image for diffusion MR images, the first inversion time image for perfusion MR images and the first TSL image for CESL images. The transformation was then applied to the atlas to yield segmentations of the brain regions co-registered to ADC and CBF maps. For perfusion MRI, images were distorted due to shimming artefacts of the EPI sequence and a nonlinear registration was used. For ADC and CESL images, affine (12 degrees of freedom) registrations were sufficient.

**Fig. 2:**
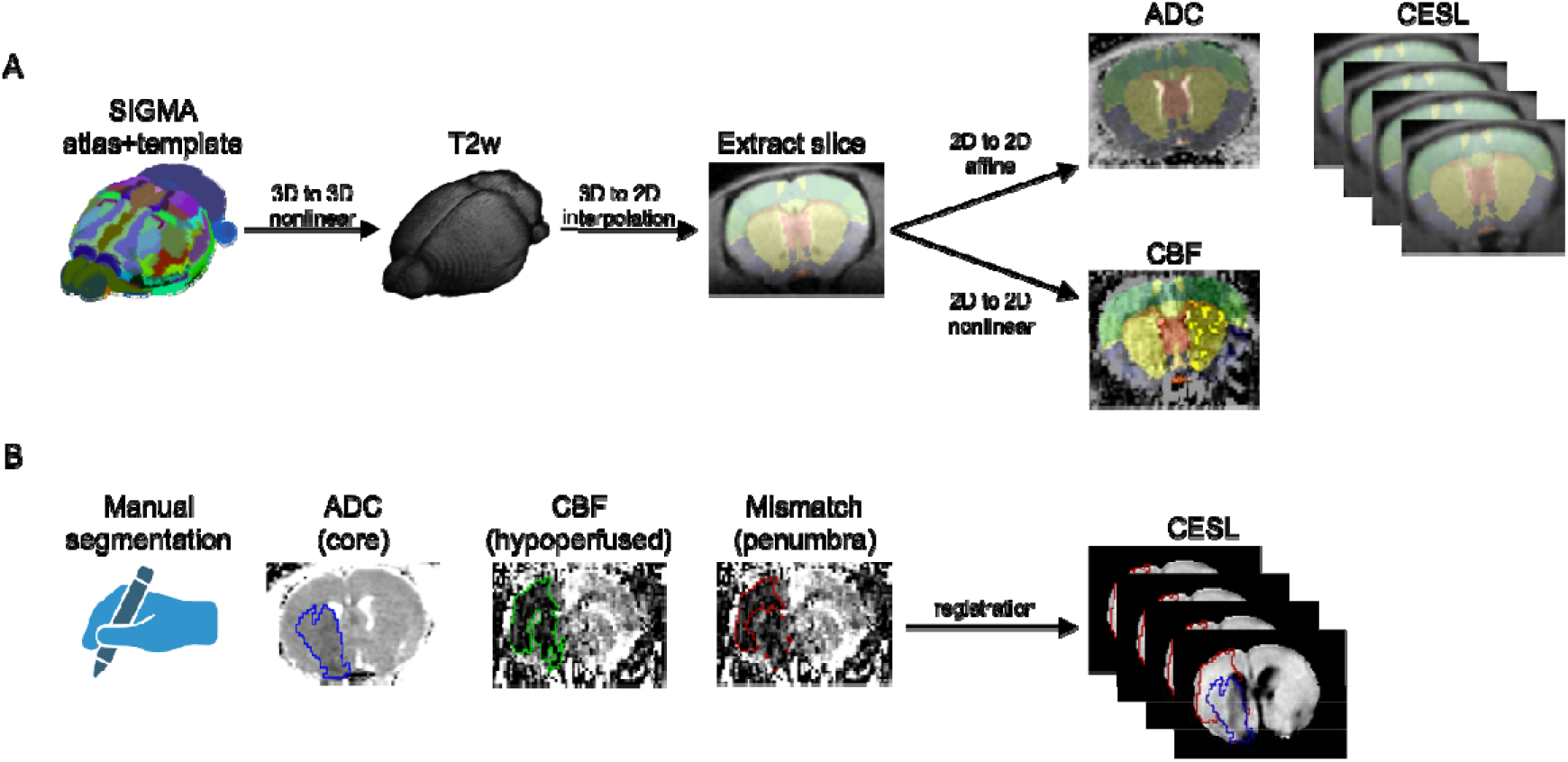
ROI analyses. CESL images were quantified in anatomical regions defined in the SIGMA rat brain atlas (A) or using manual delineation of the lesion on ADC, CBF and late ΔR1ρ maps (mean of the last 5 maps). Penumbra was defined via perfusion/diffusion mismatch.

For the second approach, the hypointense lesion was manually segmented on ADC, CBF and ΔR1ρ maps (mean image of the last 5 images) by an experienced researcher. Using the image registration transforms of the first approach, the lesion masks were transformed from ADC and CBF to the atlas and then registered to the ΔR1ρ maps. In atlas space, ROIs mirrored at the midline were automatically generated and transformed to the ΔR1ρ maps. Lesion core was defined via hypointense ADC, hypoperfused tissue via hypointense CBF and the metabolic lesion was defined via hypointense areas on the mean ΔR1ρ maps (mean of the last 5 measurements). The penumbra was defined via perfusion diffusion mismatch, i.e.

### Statistics

Statistical analyses were performed in MATLAB (version 2021a, Natick, MA, USA). Values are expressed as mean ± 95% confidence interval (CI) for time courses and the correlation analyses. For the bar plots, mean ± standard deviation is shown. Effect sizes of ipsi-vs. contralateral mean values are reported as Cohen’s d. Statistical analysis of ipsi-vs. contralateral values in the bar plots was performed using paired t-tests. Correlations of 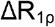 with 2DG concentration and 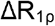 with ADC and CBF were performed using Pearson correlation analyses. Lesion masks from the different modalities were compared using mask area and using Dice similarity coefficient with the metabolic lesion mask (derived from 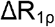 maps) with all other masks. A test with p < 0.05 was considered statistically significant. Bonferroni correction of the significance level for multiple comparisons was used where indicated in the results.

## Results

CESL MRI of phantoms containing different concentrations c_2DG_ of 2DG in buffer revealed the expected linear relationship with concentration (Fig. 3):

**Fig. 3:**
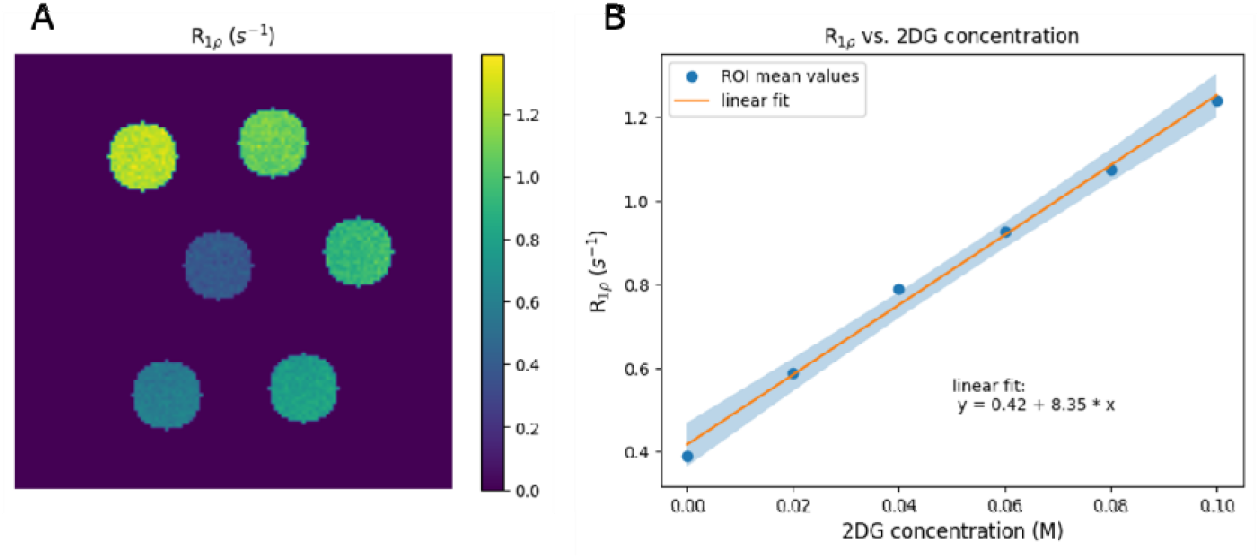
Phantom experiments. R1ρ map of NMR tubes with different concentrations of 2DG (A) and linear fit of mean R1ρ within each tube over 2DG concentration (B).

In the rat MCAO model, we found a continuous increase of 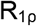 over ∼45 min CESL imaging time in metabolically active contralateral tissue whereas 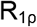 remained stable or even decreased in ipsilateral tissue (Fig. 4A, B). The decrease of R1ρ in the lesion can be explained by development of vasogenic edema and the accompanying decrease in transverse relaxation R2, which directly impacts R1ρ via R1ρ=Rex+R2. Rex summarizes relaxation in the rotating frame due to chemical exchange. When quantifying late 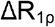 (mean of last 5 scans, Fig. 4C), the contrast was most pronounced in striatum (ipsi: - 1.54±2.62%, contra: 2.80±1.45%, t = -5.27, p = 0.00026, significant after Bonferroni correction, Cohen’s d = 1.52) and lesion core (ipsi: -1.11±2.70%, contra: 3.09±1.10%, t = - 5.08, p = 0.00036, significant after Bonferroni correction, Cohen’s d = 1.47) but smaller in hypoperfused tissue (ipsi: -1.04±2.76%, contra: 2.33±1.73%, t = -3.84, p = 0.0027, significant after Bonferroni correction, Cohen’s d = 1.11) and not significant in the penumbra (ipsi: 0.04±3.19%, contra: 0.99±2.55%, t = -1.16, p = 0.27, Cohen’s d=0.33).

**Fig. 4:**
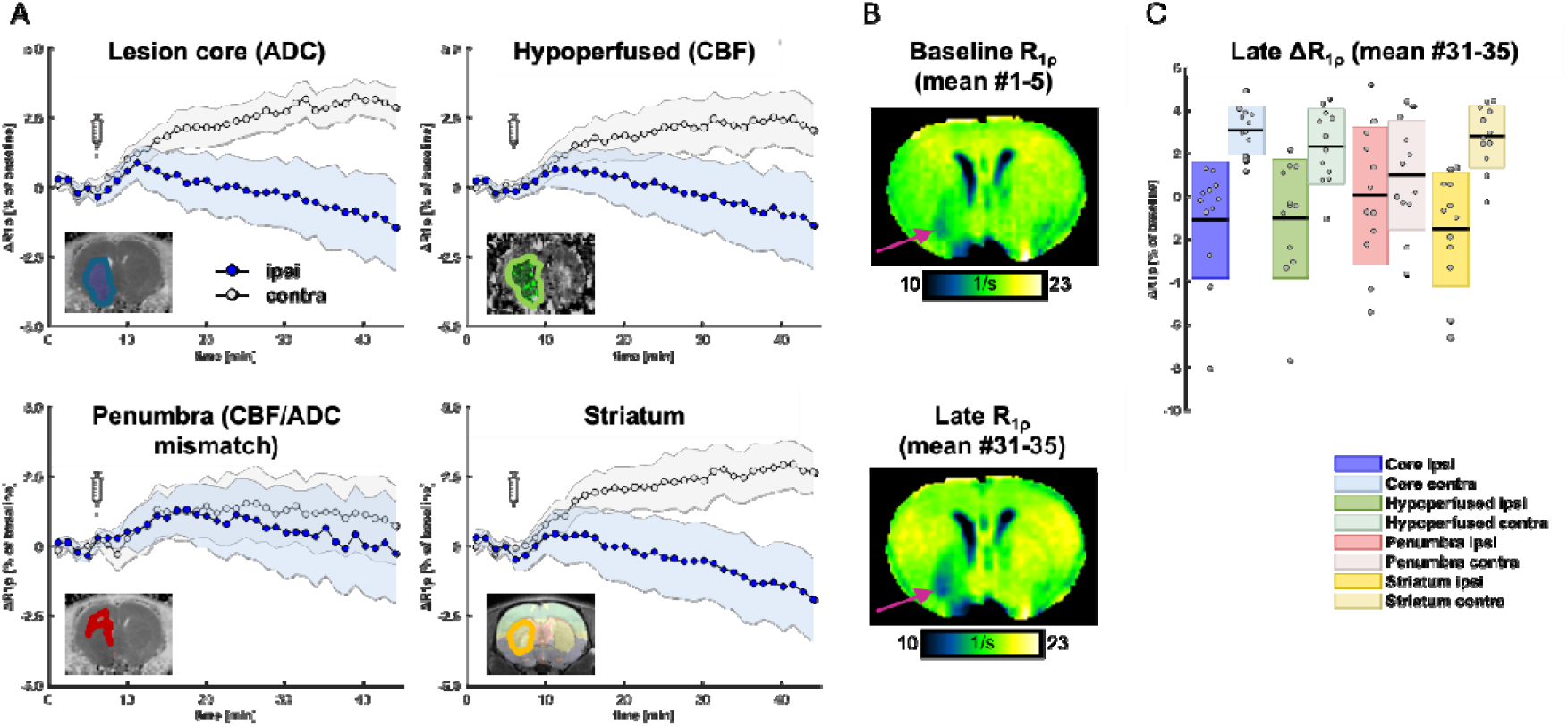
In vivo metabolic MRI. Quantification of mean 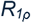 in lesion core, hypoperfused area, penumbra, ipsilateral striatum (blue) and in corresponding mirrored ROIs (gray) showed strong differences that continuously increased over time. Shaded area corresponds to 95% CI (A). Representative baseline and late (mean of first and last 5 maps) 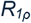 maps show an increase in contralateral tissue and slight decrease in the lesion territory (B). Quantification of late 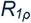 showed strongest effects of ipsi-vs. contralateral values in striatum and lesion core (C).

We then investigated how the new biomarker 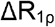 (mean of last 5 scans) correlates with conventional MR biomarkers of focal cerebral ischemia, i.e. ADC (Fig. 5) and CBF (Fig. 6). A significant correlation was found only for ADC in the ipsilateral striatum (Pearson R = 0.65, p = 0.023).

**Fig. 5:**
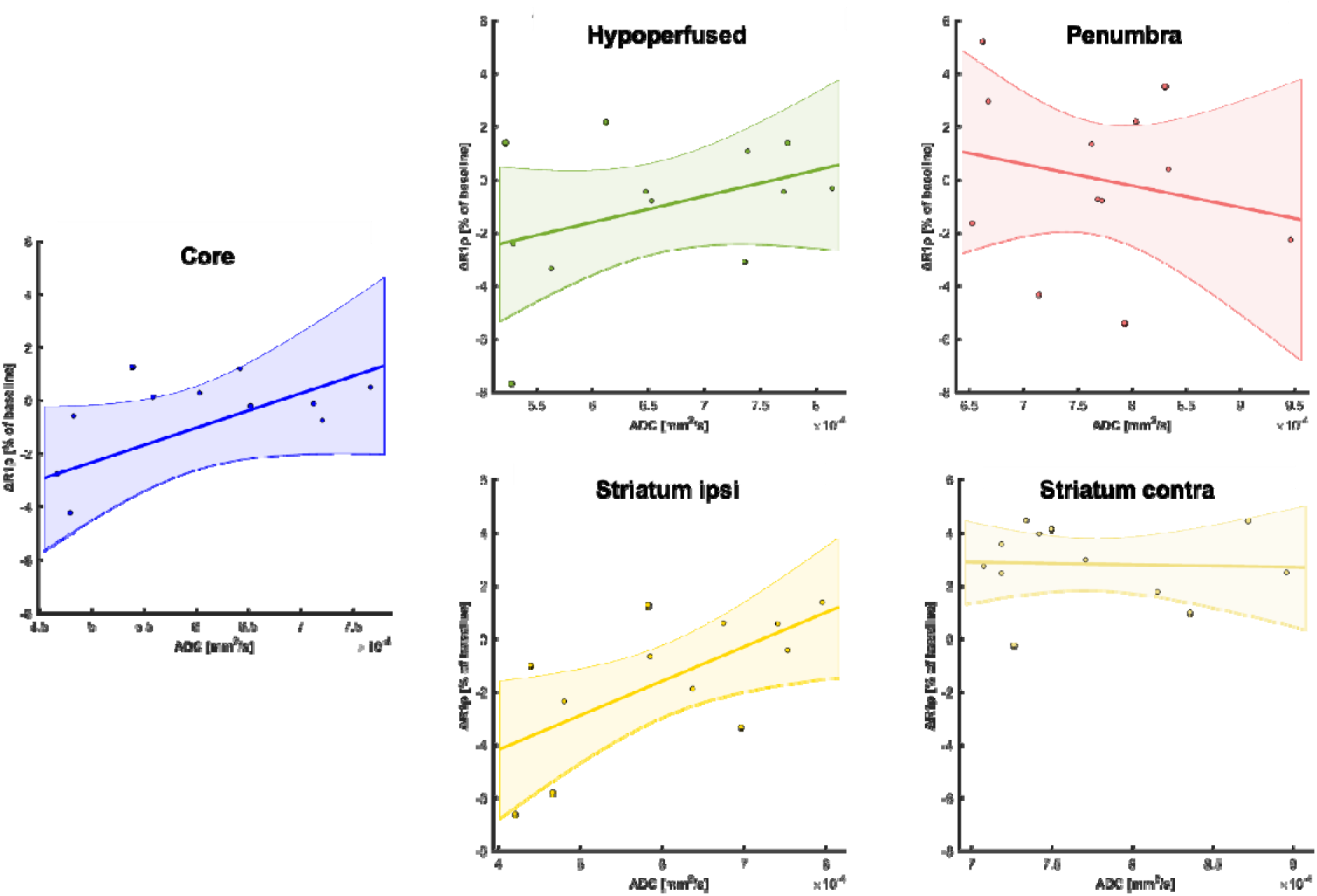
Correlation of mean late 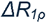; with mean ADC in ROIs.

**Fig. 6:**
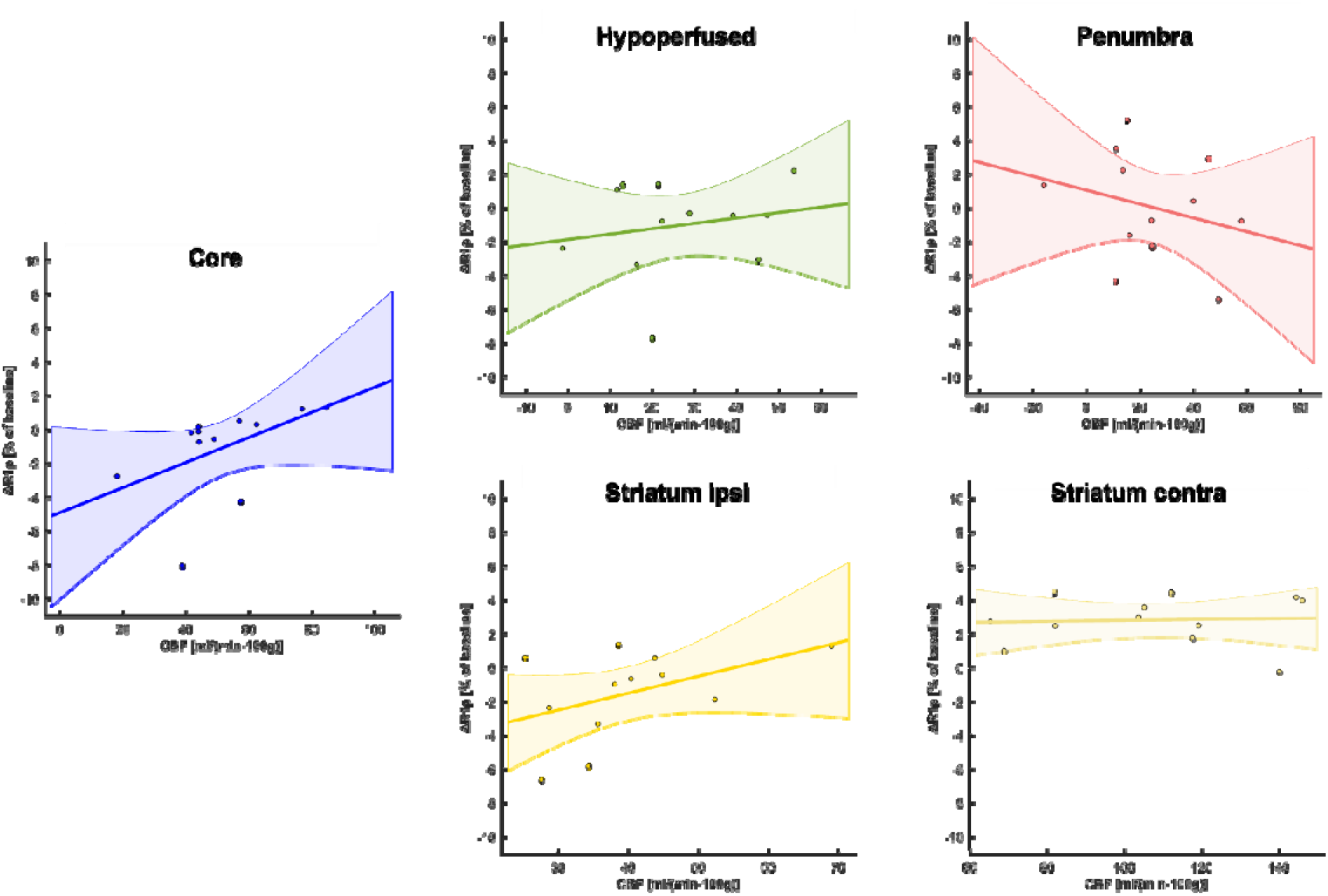
Correlation of mean late 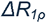; with CBF in ROIs.

Finally, in a radiologist’s approach, we wanted to investigate how the lesion mask derived from 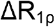 maps, i.e. the metabolic lesion, correlates with the lesion defined via ADC and CBF maps. The metabolic lesion area (Fig. 7A) was smaller than lesion core (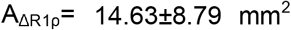 vs. A_ADC_ = 21.95±6.95 mm^2^, t = -4.94, p = 0.0004, significant after Bonferroni correction) and hypoperfused area (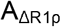 vs. A_CBF_ = 26.48± 5.76 mm^2^, t = -4.64, p = 0.0007, significant after Bonferroni correction) and comparable to penumbra (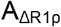 vs. A_penumbra_= 10.86± 4.81 mm^2^ t = 1.10, p = 0.30). Dice coefficient D similarity analysis (Fig. 7B) revealed most overlap with the lesion core 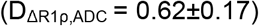, defined via ADC followed by hypoperfused area 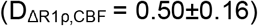 and penumbra 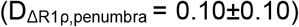. The pairwise differences between the three Dice coefficients were all significant after Bonferroni correction (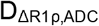 vs. 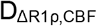, t = 5.34, p = 0.0002; 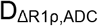 vs. 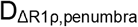, t = 8.79, p = 3*10^-6^; 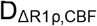 vs. 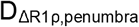, t = 8.12, p = 6*10^-6^).

**Fig. 7:**
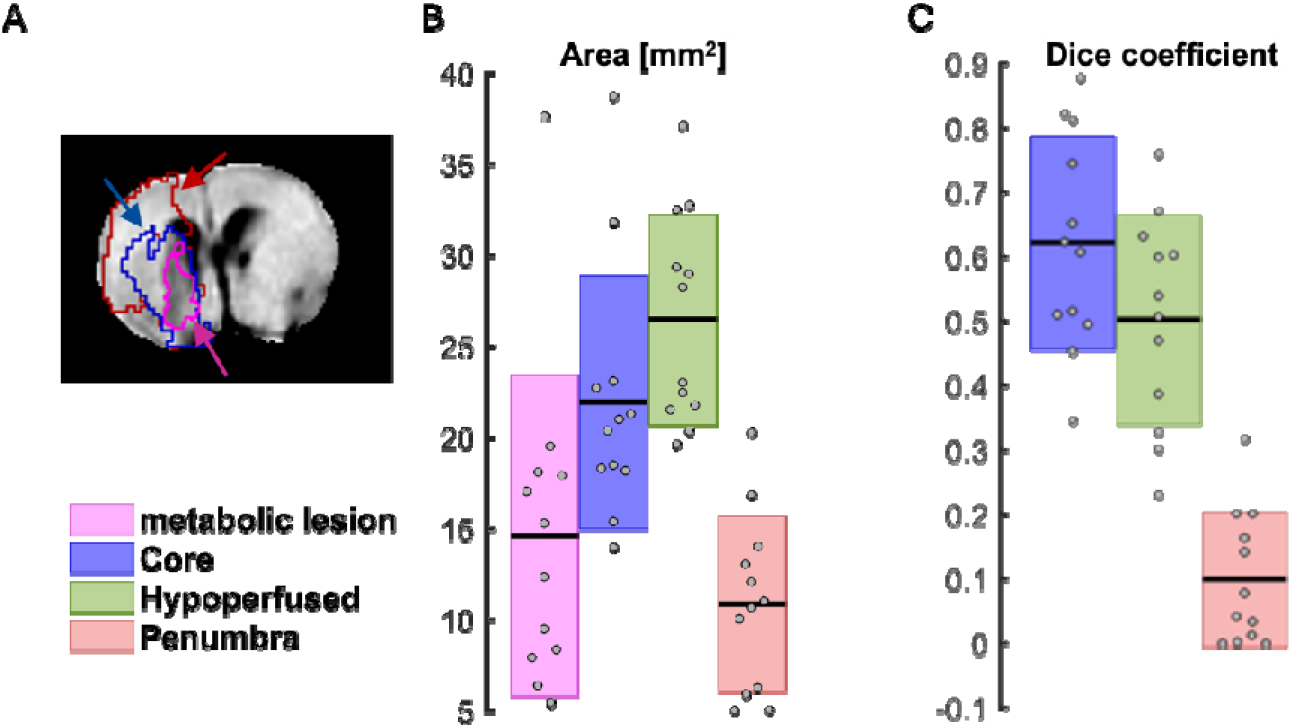
Characterization of metabolic lesion mask defined via hypointense late 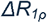;. Delineated masks overlaid on 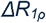; map from representative animal (A). The metabolic lesion (magenta) is smaller than lesion core (blue) or hypoperfused area (green, B) and overlaps best with lesion core compared to hypoperfused area or penumbra (red) as assessed by Dice coefficient analysis (C).

## Discussion

In this study, we established 2DG CESL MRI as a new biomarker for cerebral glucose metabolism in a rat model of ischemic stroke. We found that 2DG CESL MRI accurately delineated the ischemic core and allowed measuring cellular uptake of the glucose analogue in the penumbra. While 2DG CESL MRI provides conceptual information similar to 18F-FDG-PET imaging, the latter has not found wide adaptation due to the high costs and challenging logistics of producing radioisotopes, and the limited clinical availability of PET scanners for routine imaging. In our hands, 2DG CESL MRI allowed metabolic imaging with high effect sizes and SNR and superior spatial resolution compared to typical resolutions achieved with PET or deuterium metabolic MRI.^31^

The size of the ischemic core in metabolic MRI was substantially smaller than resolved with conventional blood flow imaging biomarkers of stroke in the present study, as expected from metabolic PET imaging.^15^ While our study was not designed to address that, this raises the question whether the metabolic lesion represents the real ischemic core. Indeed, previous work has discussed and measured the proportion by which the surrogate identification of the penumbra using flow-based MRI sequences overestimate the size of the ischemic core.^15^ Therefore, our study provides a path for establishing metabolic MRI as a precise tool to measure the penumbra. A previous study investigating the utility in early MRI in predicting final infarct sizes in murine stroke demonstrated that the accuracy of ADC to predict the final infarct size depends on the time of imaging.^32^ It is thus conceivable that the timing of imaging in our study was can explain differences and that the metabolic infarct core measured early after reperfusion is not representative of the final infarct size.

As acute treatments like endovascular thrombectomy broaden therapeutic options for stroke, advanced imaging modalities play a crucial role in identifying patients who may benefit most from interventional therapy. The mismatch between perfusion and diffusion deficits is often considered a robust indicator of salvageable brain tissue,^33, 34^ and mismatch imaging has become a crucial tool for stratifying patients and selecting therapies.^34, 35^ However, the clinical value and accuracy of the perfusion/diffusion mismatch in defining the penumbra is still under debate,^15, 33^ which calls for further research into alternative and more precise methods. To this end, integrating metabolic imaging, and precisely quantifying both the metabolic stroke core and metabolic penumbra at the time of emergency care, could further refine these treatment decisions. By providing essential data on tissue viability, metabolic imaging can improve our understanding of salvageable brain regions and inform on the potential reversibility of clinical deficits.

It should be noted that due to the high concentration of 2DG required for CESL MRI compared to FDG PET, it is presently not suitable as a translational tool. However, a previous study has used regular D-glucose for cerebral CESL imaging with reasonable signal-to-background,^21^ albeit different temporal kinetics, indicating that alternative carbohydrate tracers could provide a viable pathway for future clinical translation with further optimization. It was shown that after injection of D-glucose or glucose analogs, 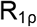 increased linearly with local concentration of the metabolite.^20, 21, 36^ The authors concluded that 2DG CESL MRI can measure glucose transport and metabolism noninvasively in normal and pathological brain tissue. However, 2DG is trapped in cells and not further metabolized. 2DG CESL MRI is limited in that regard in the same way as 18F-FDG-PET. To this end, the recent introduction of dynamic nuclear polarization (hyperpolarization) has made it possible to map the metabolism of carbon-13-labeled metabolic substrates in preclinical models^37, 38^ and in humans^38-40^ and achieves high spatial and temporal resolution as well as substantially improved signal-to-noise ratios.^38^ Another recent study used deuterium (2H) metabolic imaging in combination with FDG-PET to study cerebral glucose metabolism after MCAO in mice.^31^ While this approach allows for the mapping of glucose metabolites, it is at the expense of spatial resolution.^31^

In summary, our present study demonstrates that 2DG CESL MRI provides new imaging biomarkers in preclinical stroke models which allows for multiplexing with other MR imaging modalities.

## Acknowledgements

We thank Dr. Kai Herz for providing the adiabatic HSExp tipping pulses and Dr. Steffen Görke and Dr. Philip Boyd for helpful discussions during the planning of the study.

## Author contributions

*Concept and design:* Boehm-Sturm, Mergenthaler

*Acquisition, analysis, or interpretation of data:* Boehm-Sturm, Schuenke, Foddis, Mueller, Koch, Mergenthaler

*Discussion of the data:* Boehm-Sturm, Schuenke, Foddis, Müller, Koch, Beard, Holloway, Mottahedin, Schröder, Buchan, Mergenthaler

*Drafting of the manuscript:* Boehm-Sturm, Mergenthaler

*Critical review of the manuscript for important intellectual content:* Boehm-Sturm, Schuenke, Foddis, Mueller, Koch, Beard, Holloway, Mottahedin, Schröder, Buchan, Mergenthaler

*Statistical analysis:* Boehm-Sturm, Schuenke, Koch

*Supervision:* Boehm-Sturm, Mergenthaler

## Statements and declarations

### Ethical considerations

Not applicable.

### Consent to participate

Not applicable.

### Consent for publication

Not applicable.

### Declaration of conflicting interest

AMB is cofounder of BRAINOMIX. DJB is an inventor, and The University of Newcastle is applicant for the patent: *Stroke Treatment (WO 2022/072348)*. The other authors report no conflicts.

### Funding statement

This work was funded by the Einstein Foundation Berlin (EJF-2020-602 to PM, EVF-2021-619 and EVF-2021-619-2 to PM and AMB) and the Leducq Foundation for Cardiovascular and Neurovascular Research (Leducq Foundation Trans-Atlantic Network of Excellence on Circadian Effects in Stroke, 21CVD04, PM and AMB). Funding to SM, MF, SPK and PBS was provided by the German Federal Ministry of Education and Research (BMBF) under the ERA-NET NEURON scheme (01EW2305), and the German Research Foundation (DFG, project BO 4484/2-1, Project-ID 424778381-TRR 295 ReTune and EXC-2049-390688087 NeuroCure). Noninvasive imaging experiments were supported by Charité 3R – Replace | Reduce | Refine. Funding to PS was provided by the Deutsche Forschungsgemeinschaft (DFG, Project-ID 372486779-SFB 1340 Matrix in Vision). DJB was supported by an EMBO Short-term Fellowship and funding from the National Health and Medical Research Council Australia (APP1182153). PM is Einstein Junior Fellow and AMB is Einstein Visiting Fellow, both funded by the Einstein Foundation Berlin. PM acknowledges funding from the Einstein Foundation Berlin (EJF-2020-602, EVF-2021-619, EVF-2021-619-2, EVF-BUA-2022-694), the Volkswagen Foundation (9A866), the Else Kröner-Fresenius Stiftung (2019-A34), and the Stiftung Charité (StC-VF-2024-59). Besides funding, the sponsoring organizations did not play any role in the preparation, review, or approval of the article, or decision to submit the article for publication.

### Data and code availability

Data and code associated with this manuscript is available on Zenodo (https://zenodo.org/records/14526092; DOI: 10.5281/zenodo.14526091).

## Notes

https://zenodo.org/records/14526092

### Summary of Updates

Methods and discussion sections have been edited for clarification. Figures and manuscript text have been edited for clarification throughout.

